# C1 compounds shape the microbial community of an abandoned century-old oil exploration well

**DOI:** 10.1101/2020.09.01.278820

**Authors:** Diego Rojas-Gätjens, Paola Fuentes-Schweizer, Keilor Rojas-Jimenez, Danilo Pérez-Pantoja, Roberto Avendaño, Randall Alpízar, Carolina Coronado-Ruíz, Max Chavarría

## Abstract

The search for microorganisms that degrade hydrocarbons is highly relevant because it enables the bioremediation of these substances cheaply and without dangerous by-products. In this work, we studied the microbial communities of an exploratory oil well, abandoned a century ago, located in the Cahuita National Park of Costa Rica. Cahuita well is characterized by a continuous efflux of methane and the presence of a mixture of hydrocarbons including C2-dibenzothiophene, phenanthrene or anthracene, fluoranthene pyrene, dibenzothiophene, tricyclic terpanes, pyrene, sesquiterpenes, sterane and n-alkanes. Based on the analysis of 16S rRNA gene amplicons, we detected a significant abundance of methylotrophic bacteria (*Methylobacillus* (6.3-26.0 % of total reads) and *Methylococcus* (4.1-30.6 %)) and the presence of common genera associated with hydrocarbon degradation, such as *Comamonas* (0.8-4.6 %), *Hydrogenophaga* (1.5-3.3 %) *Rhodobacter* (1.0-4.9 %) and *Flavobacterium* (1.1-6.5 %). We evidenced the presence of methane monooxygenase (MMO) activities, responsible for the first step in methane metabolism, by amplifying the *pmo* gene from environmental DNA. We also isolated a strain of *Methylorubrum rhodesianum*, which was capable of using methanol as its sole carbon source. This work represents a contribution to the understanding of the ecology of communities of microorganisms in environments with permanently high concentrations of methane and hydrocarbons, which also has biotechnological implications for the bioremediation of highly polluting petroleum components.

## Introduction

Hydrocarbon pollutants have been declared an environmental issue [1], as a result of their slow biodegradability, genotoxic, mutagenic and cytotoxic activities, especially polycyclic aromatic hydrocarbons (PAH), as well as saturated and unsaturated aliphatic hydrocarbons [2–3]. Hydrocarbon-degrading bacteria have gained importance in recent decades due to their potential in bioremediation and their metabolic versatility. Several studies have been undertaken to understand the mechanisms, paths and composition of bacterial communities involved in the microbial degradation of hydrocarbons [4–7]. Numerous strains with the ability to use these compounds have been isolated and studied; prominent examples are *Pseudomonas putida* [8–9], *Bacillus* sp. [10–11], *Burkholderia* sp. [12–13], *Sphingomonas* sp. [14] among others [15–17]. Even though isolated strains show a fascinating potential under controlled conditions [18–19] most of them failed in field experiments *in situ* [20–22]. Moreover, culture-dependent techniques (i.e. a study with bacterial isolates) are unable to completely explain the ecology of their habitats. Novel efforts are thus focused on hydrocarbon-degrading bacteria consortia. One way to obtain a general overview of the microbial consortia responsible for degrading hydrocarbons is through culture-independent techniques. Studies with culture-independent techniques have been developed in waters [23–24], soils [25–27], and deep-sea sediments [16,28–29] contaminated with hydrocarbons. Experiments with controlled environments supplemented with multiple oils or biosurfactants [30] have also been conducted. Although these studies are interesting, they have been conducted in environments in which the contact with hydrocarbons was for a brief duration and do not represent accurately the conditions that occur in an oil well in which hydrocarbons are mixed with other carbon sources and gases [31–32]. Few studies have been performed on oil wells; most of them evaluate only the biodegradation capabilities or utilize culture-dependent methods [33–34]. Other studies of oil wells apply to extreme environments with high pressure and temperature, reflecting a community mainly modeled by these physical conditions rather than by the carbon source [35–36]. Therefore, the study of environments where a microbial community is permanently confronted with hydrocarbons and not transiently as occurs in oil spills, is a topic of great interest to understand the ecology and evolution of microorganisms in sites with high concentrations of hydrocarbons as well as for future biotechnological applications.

In this work, based on the analysis of 16S rRNA gene amplicons, we performed an analysis of the microbial communities in an abandoned oil exploration well that dates back almost a century. This oil well is located in Cahuita National Park (for which reason the well is known as Cahuita N° 1; Figs. 1a and 1b) in Limón, Costa Rica, which was drilled in 1921 by Sinclair Central American Corporation [37]. This oil well is emblematic in Costa Rican history as it represents an early attempt to obtain oil in the country. After a few years of operation, the efforts to obtain oil ceased; the exploratory oil well was abandoned and has currently become a touristic attraction in a protected zone. Nevertheless, the well was sealed improperly and still expels hydrocarbons and gases to the environment (see Fig. 1c). The few existing records indicate that Cahuita well N° 1 has a depth between 1777 [37] and 1922 m [38] but a complete physicochemical characterization has not be performed yet. The persistence of hydrocarbons and gases in the well for almost 100 years makes it an attractive candidate for an analysis of the microbial composition to understand which organisms are subsisting in a long-term hydrocarbon-providing environment.

**Fig 1.**
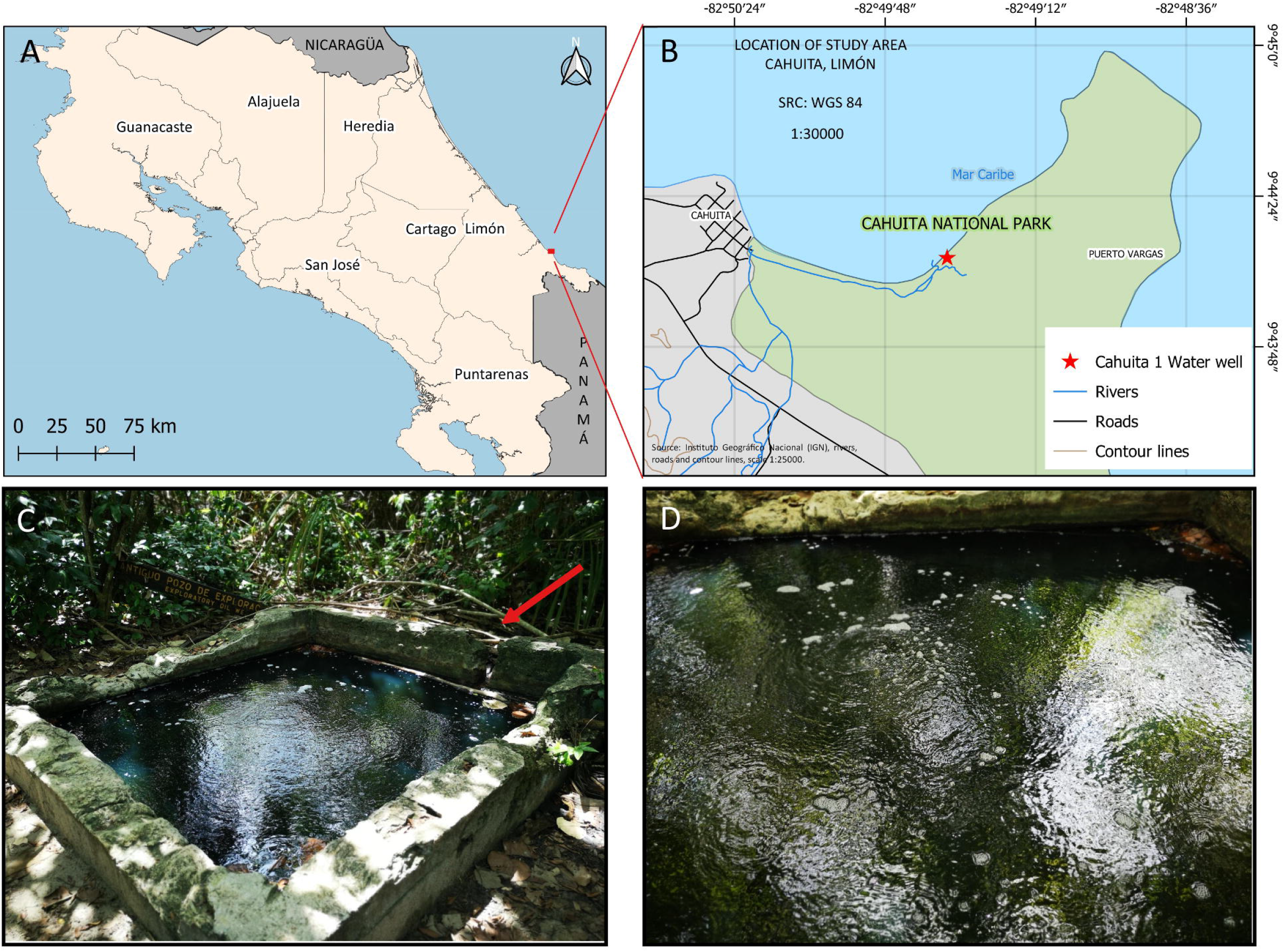
Exploratory oil well at Cahuita, Limón, Costa Rica. A) This exploratory oil well is located at the Caribbean coast of Costa Rica in Cahuita National Park. B) The exploratory oil well is located beside Suarez River. The red star shows the exact location. C) The well was drilled in 1921; a cement base about 3 × 3 m was built. As seen in the figure, the well has an oily appearance and is slightly darker than normal waters. The red arrow shows where contaminated water drains to the soil and then to the river. The sediments were taken from the contaminated soils. D) From the surface of the well a constant efflux of gas occurs.

## Materials and Methods

### Study Site

The old oil exploration well (latitude 9.7359°, longitude −82.8259°) is located in Cahuita National Park (Figs. 1a and 1b) in Limón, Costa Rica. The well has a square cement structure with surface area ~9 m^2^ (Fig. 1c), depth 2.20 m and is full of water of clear blackish hue. The center of the well that exhibits a continuous bubbling of gas (Fig. 1d, see suppl. video S1) is presumably a narrow hole of depth 1777-1922 m reported in the literature (Castillo, 1961; Pizarro A, 1993). All permissions required for sampling the exploratory oil well were obtained from National System of Conservation Areas (SINAC), Ministry of Environment and Energy (MINAE) of Costa Rica (resolutions R-SINAC-PNI-ACLAC-013-2019, R-SINAC-PNI-ACLAC-007-2020 and R-SINAC-PNI-ACLAC-022-2020) and Commission of Biodiversity of University of Costa Rica (resolution N° 147).

### Sampling and field measurements

Three sampling campaigns were conducted in this work. In the first campaign (2019 June), nine water samples (each 1 L) were collected in sterile glass bottles at various points on the surface of the 9 m^2^ well. Those nine samples were pooled into three composite samples (codes SW1, SW2, SW3; SW means surface water). In the second campaign (2019 August), nine samples (each 1 L) were taken from the center of the well along the depth of the cement structure, specifically three water samples from the surface (pooled and named SW4), three water samples at depth 1 m (pooled and named W1M; i.e. water 1 m) and three water samples at depth 2.20 m (pooled and named W2M; i.e. water 2 m). For analysis of 16S rRNA gene amplicons, some samples were processed in duplicate, so indicated in the name R1 (replica 1) or R2 (replica 2). Multiple attempts were made to introduce a probe into the main borehole (i.e. the hole of depth 1777-1922 m), but it seems that, over time, this narrow hole has become obstructed by sediments and various materials, which allow only the exit of gases or liquids. In the field, the temperature and dissolved oxygen (DO) were measured in water samples with a dissolved-oxygen meter (Model 85, Yellow Springs Instrument Company Inc, Ohio, USA). Water samples for both chemical and microbial analysis were collected using sterile glass bottles and were immediately transported to the laboratory and stored at 4 °C until processing. For sediment sampling, masses between 10 and 50 g were placed in sterile centrifuge tubes (50 mL). Sediments (SED1 and SED2) were gathered in the area indicated in Fig. 1c, a few cm after the well drain to the estuary. The sediments were collected at depths not exceeding 15 cm. Gas samples were collected into a bag (Flexfoil®, cat 262-01, SKC, USA) assisted with a plastic funnel and a grab air-bag sample pump (cat 222-2301, SKC, USA). In the third campaign (July 2020) five samples from the superficial water were taken specifically for microbial isolation.

### Measurements of physicochemical parameters

The content of calcium, iron, magnesium, potassium, sodium, phosphorus, alkalinity, and total sulfur was analyzed for all samples following Standard Methods for the Examination of Water and Wastewater (SMWW) described by APHA (2017) [39]. The total and partial alkalinity were measured according to APHA (2017) (method 2320B) [39]. The pH of the water samples was measured with a pH meter (Mettler Toledo Seven compact duo S213, Columbus, Ohio, USA). The conductivity of the samples was measured with a conductivity meter (Mettler Toledo Seven compact duo S213, Columbus, Ohio, USA). The probe was calibrated with a standard solution of known conductivity. The metals were analysed based on APHA standards (method 2320B, Waltham, MA, USA) with atomic absorption (ICP-OES, Perkin Elmer, model Optima 8300, MA, USA). Spectrophotometric determination of ammonium, nitrate and ureic nitrogen concentrations were made in triplicate (10 mL per sample) following QuickChem® methods 10-107-04-1-A, 10-107-06-2-O and 31-206-00-1-A respectively in a flow-injection analyzer (FIA, LACHAT QuickChem 8500 Series 2, Mill Rd, WI, USA). Before samples in each batch were analyzed, calibration curves were prepared for each analyte. All analyses were performed at Agronomical Research Center – University of Costa Rica (CIA-UCR) and Center for Electrochemistry and Chemical Energy (CELEQ-UCR).

### Analysis of hydrocarbons and gases

The extraction and qualitative determination of hydrocarbons were performed according to the method described by the European Normalization Committee (2009) [40]. Liquid-liquid extraction (sample 20 mL) with three portions of dichloromethane (each 20 mL) was performed. The extract was eluted with a funnel and sodium sulfate and then concentrated using a rotary evaporator initially, followed by a flow of nitrogen to volume 500 μL. From that volume, 50 μL was extracted and eluted in a second alumina column previously deactivated with hexane (7 mL). A gas chromatograph (Agilent Technologies, model 7890B, CA, USA) with a capillary column (Agilent Technologies, HP-5-MS, 30 m × 0.25 mm × 0.25 μm, CA, USA), coupled to a mass spectrometer (MS, Agilent Technologies, model 5977, CA, USA) with helium UAP carrier gas (flow rate 1.1 mL/min) was used to perform the analysis. The temperature of the oven was maintained at 42 °C for 1.30 min and then a ramp (5.5 °C/min) was applied until 300 °C. The data were analyzed (MSD ChemStation program, version E.02.00.493). For the SCAN mode method, a range 50-450 u was employed. Methanol was measured with a gas chromatograph (Agilent Technologies, model 7890B, CA, USA) with an analytical column (DB-WAX, Agilent Technologies, 30 m × 0.25 mm ID × 0.25 μm) and measured simultaneously with both a flame-ionization (FID) and MS detector. The chromatograph was operated with a temperature ramp (50 °C for 5 min and then at 25 °C/min to 160 °C); the detector and the injector were set at 280 °C. Argon served as carrier gas (4.0 mL/min). Highly pure standards were used (methanol Sigma Aldrich CAS 180 67-56-1). The composition of the gas (specifically the content of methane, ethane, nitrogen, oxygen, carbon oxide and carbon dioxide) was measured with a gas chromatograph (Hewlett Pachard, 6890, CA, USA) equipped with a PLOT fused silica column (GS-Gaspro, Agilent Technologies, 30 m × 0.32 mm) and both thermal-conductivity (TCD) and FI detectors. The chromatograph was operated with a temperature ramp (from 60 °C for 6 min and then at 25 °C/min to 260 °C). The injector and detector temperatures were both 250 °C; argon served as carrier gas (4.5 mL/min). Certified standard gas (Agilent Technologies, type refinery gas, capacity 1 L, pressure 30 psig at 21 °C, batch number 112PLU1SPC10D, CA, USA) served as a standard for accuracy. All analyses were performed at CELEQ-UCR.

### DNA Extraction and 16S rRNA gene sequencing

Samples were processed as described previously [41] with some modifications. Briefly, each sample was filtered individually through a vacuum system under sterile conditions using a sterile membrane filter (pore size 0.22 μm, GV CAT No GVWP04700, Millipore, Darmstadt, Germany). The filters were collected and stored at −30 °C until processing, and then cut into small pieces using sterile surgical scissors. The DNA was extracted with a DNA isolation kit (PowerSoil®, MoBio, Carlsbad, CA, USA) as described by the manufacturer; half a filter was used for each extraction. Cell lysis was accomplished in one step of bead beating (FastPrep-24, MP Biomedicals, Santa Ana, CA, USA) for 40 s at 6.0 m/s. To process the sediments, a representative sample of mass 500 mg (obtained with vortex homogenization and then through grid sampling) was collected from 10-50 g of each wet sediment; DNA was extracted according to the same protocol. For the construction of the microbial amplicon library (16S rRNA), the V4 hypervariable region was PCR-amplified with universal primers 515F and 806R [42]. The PCR-generated products were subjected to a 250 nt paired-end sequencing (Illumina MiSeq, Novogene Bioinformatics Technology Co., Ltd, CA, USA).

### Bioinformatic analysis of 16S rDNA amplicon data

Sequences were quality check and analyzed using version 1.12 of the DADA2 pipeline [43]. Briefly, reads were quality-filtered, deduplicate and Amplicon Sequence Variant (ASV), a higher-resolution analogue of the traditional Operational Taxonomic Unit (OTU) were inferred. Finally, chimeric sequences and global singletons were removed. This process resulted in an amplicon sequence variant (ASV) table, which records the number of times each exact amplicon sequence variant was observed in a sample. The taxonomic assignment was performed on comparing sequences against the SILVA reference database v132 [43] and then curated on comparing sequences against the GenBank database of NCBI. Global singletons were removed. After these processes, we obtained 2,561,563 sequences and 6993 ASV. The average number of sequences per sample was 197,043, ranging from 181,657 to 225,244. The sequence data were deposited in the sequence-read archive (SRA) of GenBank under the BioProject ID PRJNA614582. The statistical analyses and their visualization were performed with the R statistical program [45] and Rstudio interface. Package Vegan v2.5-6 [46] was used to calculate alpha-diversity estimators (Shannon and Simpson indexes), non-metric multi-dimensional scaling analyses (NMDS). Data tables with the ASV abundances were normalized into relative abundances and then converted into a Bray–Curtis similarity matrix. The differences of the oxygen concentration (See Table 1) on the bacterial community composition were evaluated with non-parametric multivariate analysis of variance (PERMANOVA) and pairwise PERMANOVA (adonis2 function with 999 permutations), taking into consideration 3 groups: superficial (0 meters), subsuperficial (1-2 meters) and sediments.

**Table 1.** Physical properties and chemical composition of the abandoned exploratory oil well. ND: Not detected. NM: Not measured.

| Parameter/element | SW1 | SW2 | SW3 | SW4 | W1M | W2M |
| --- | --- | --- | --- | --- | --- | --- |
| pH ($\pm 0.1$) | 7.5 | 7.4 | 6.9 | 7.9 | 8.0 | 8.0 |
| Oxygen /% $\pm 0.01$ | NM | NM | NM | 12.10 | 13.20 | 6.76 |
| Ammonium /mg L<sup>-1</sup> $\pm 0.1$ | 2.0 | 2.0 | 1.4 | 2.4 | 2.4 | 2.4 |
| Nitrate /mg L<sup>-1</sup> $\pm 0.005$ | 0.100 | 0.200 | 0.200 | 0.100 | 0.100 | 0.100 |
| Ureic nitrogen/mg L<sup>-1</sup> $\pm 0.005$ | 0.100 | 0.100 | 0.100 | 0.100 | 0.100 | 0.100 |
| Calcium /mg L<sup>-1</sup> $\pm 2$ | 42 | 43 | 41 | 26 | 25 | 26 |
| Magnesium /mg L<sup>-1</sup> $\pm 2$ | 17 | 16 | 16 | 10 | 10 | 11 |
| Potassium /mg L<sup>-1</sup> $\pm 2$ | 19 | 24 | 22 | 7 | 8 | 8 |
| Phosphorus /mg L<sup>-1</sup> $\pm 0.01$ | ND | ND | ND | 0.10 | ND | ND |
| Iron /mg L<sup>-1</sup> ± 0.01 | ND | ND | ND | 0.10 | 0.10 | 0.10 |
| Sodium /mg L<sup>-1</sup> ± 40 | NM | NM | NM | 780 | 788 | 781 |
| Sulfur /mg L<sup>-1</sup> ± 0.05 | 0.70 | 0.30 | 0.40 | 0.80 | 0.90 | 1.60 |
| Alkalinity /mg L<sup>-1</sup> ± 5 | 138 | 132 | 107 | 148 | 142 | 133 |

### Methane monooxygenase (MMO) detection

For determining the presence of the enzyme Methane Monooxygenase (MMO) a section of the gene *pmo* was amplified using primers PmoC374 (AGCARGACGGYACNTGGC) and PmoA344 (ANGTCCAHCCCCAGAAGT) described by Ghashghavi et al. [48]. Due to the degenerate nature of primers selected, the polymerase chain reaction (PCR) was performed on a Mastercycler ep Gradient S thermal cycler (Eppendorf, Hamburg, Germany) using a slow ramp in temperature (0.1 °C s^−1^) between the annealing and extension cycles, as described by Luton et al. [49]. A negative control was prepared with all components of the PCR reaction except the DNA sample. PCR products were analyzed on 2% agarose gels using standard protocols. For sequencing a unique visible band in the agarose gel of ~700 bp was excised using a sterile bistoury and purified using GeneJET Gel Extraction Kit (Thermo Scientific™, USA). Finally, PCR products were sequenced using BigDye Terminator v3.1 cycle sequencing kit and cleaned with BigDye XTerminator Purification kit (Applied Biosystems, USA). Products were analysed on a 3130xl Genetic Analyzer (Applied Biosystems, USA). Forward and Reverse sequences were assembled using Bioedit. Nucleotide sequence was deposited on the GenBank database under accession number MT948143.

### Isolation, identification, and phylogenetic analysis of methylotrophic bacteria

Water samples (1L) from the well were filter using a sterile membrane filter (pore size 0.22 μm, GV CAT No GVWP04700, Millipore, Darmstadt, Germany). The filters were collected and then resuspended in Dorn mineral medium [50] with methanol 25 mM as a sole source of carbon and energy. A dense 20 mL culture was obtained after 3 days of incubation at 30°C in fluted 100 mL Erlenmeyer flasks on a rotary shaker (180 rpm). A 20 μL aliquot of this culture was successively transferred by three times into fresh medium. Lastly, 50 μL of the culture was plated on Dorn mineral medium agar plates containing methanol as the sole carbon source (25 mM). One of the fastest growing colonies was picked and streaked on the same agar medium. Purity was checked on R2A agar (Merck KGaA, Darmstadt, Germany) plates and the ability to use methanol as sole carbon and energy source was again checked in liquid mineral medium.

For identification by 16S rRNA gene sequencing, a small inoculum of the bacteria was added to the Polymerase chain reaction (PCR) mixture. PCR and sequencing protocol were made as described above using the primers 27F and 1492R [51]. The obtained sequence was compared to the 16S rRNA bacterial/Archaeal database of the NCBI using blast. Nucleotide sequence was deposited on the GenBank under accession number MT936438. For phylogenetic analysis, the closest 16S rRNA sequences from validly described microbial type strains and isolates were retrieved by blastN against the curated 16S rRNA database. The sequences were aligned using MUSCLE [52]. These alignments enabled phylogenetic reconstruction with MEGA6 software [53] and the maximum-likelihood method based on the general time-reversible model. In total, 1000 bootstrap replications were calculated to ensure the robustness of the results.

### Bacterial growth using methanol as its sole carbon source

Growth curves were performed to evaluate the ability of the bacterial isolate to use methanol as the sole carbon source. First, bacterium was striked in solid M9 mineral medium supplemented with 25 mM of methanol as its sole carbon source. Then a preinoculum of the bacterium was growth for 48 h in M9 mineral medium with 25 mM of methanol. Later, the culture was diluted to an initial optical approximately 0.05 at 600 nm in fresh M9 mineral medium containing methanol as its sole carbon source (25 mM). Bacterial growth was estimated on monitoring the optical density at 600 nm (Synergy H1 Hybrid Multi-Mode Reader, Biotek, Winooski VT, USA) over 4 independent replicates of cultures in plates (96 wells; Nunclon 162 TM Surface; Nunc A/S, Roskilde, Denmark). Plates were incubated at 30 °C for 48 h with continuous orbital shaking at 180 rpm; the optical density was measured every 5 min.

## Results and Discussion

### Physicochemical analysis of Cahuita well N° 1

The major physical-chemical properties of the oil well samples are presented in Table 1. The data revealed that Cahuita well N° 1 presented a pH neutral to slightly alkaline (7.0-8.0) and corresponds to a dysoxic environment (0.63-0.90 mg O_2_/L, Table 1) according to terminology proposed by Tyson and Pearson (1991) [54]. The chemical analyses revealed the presence of nitrate, typically found in most fresh and sea water as well as urea-nitrogen and ammonium. The latter two chemical forms of nitrogen are associated with organic material; their presence in the oil well is reasonable in view of the reducing environment of these sites, the presence of vegetation and the influence of sea water [55]. The concentrations of calcium, magnesium, potassium, iron and sulfur are in the ranges accepted for freshwater according to the Costa Rican Institute of Water and Sewers (AyA) [56]. The concentration of sodium and the conductivity measurements indicate that fresh and sea water are present in combination in the sampled site. (see Table 1). This condition is likely due to the fact that the oil well is near the coast (Fig. 1b), and seawater seeps through the ground until it reaches the well, increasing the values of these parameters.

As mentioned above, Cahuita well N° 1 shows a constant efflux of gases rising from the center of the well (Fig. 1d, see Supplementary Video S1). GC-analysis shows that the gas is mostly methane (Supplementary Fig. S1). Other gases such as ethane, N_2_, O_2_, CO and CO_2_ were not detected. Previously, it has been reported that most oil wells are also sources of natural gas, mainly comprising methane (Alvarez et al. 2018; Zavala-Araiza et al. 2015). Other C1 compounds such as methanol were not detected (limit of detection 80 mg/L or 2.5 mM).

Various hydrocarbons were also identified in all samples (see Fig. 2). Samples taken during the first sampling (SW1, SW2, SW3) contained C2-dibenzothiophene, phenanthrene or anthracene, fluoranthene, dibenzothiophene, tricyclic terpane, pyrene, sesquiterpenes, sterane and n-alkanes. In the second sampling (samples SW4, W1M and W2M), only C2-dibenzothiophene and steranes were found. We suggest that the absence of most hydrocarbons in the second sampling could be due to high tide, which dilutes these products in the soil and the oil well. Other simple aromatic compounds commonly known as BTEX (benzene, toluene, ethylbenzene and xylene) were not identified in any sample. Nevertheless, C2-dibenzothiophene was identified in all samples, and during the low tide n-alkanes, steranes and sesquiterpenes were all present. PAH (including C2-dibenzothiophene) are important pollutants of the environment because of their extremely cytotoxic, genotoxic and mutagenic activity and slow biodegradability [57]. Aliphatic and polycyclic saturated hydrocarbons are also important contaminants of the environment because they readily diffuse in soil and subterranean water reservoirs [58]. These results indicate that the samples from an exploratory oil well contain hydrocarbons and pollutant gases of significant importance for the environment in a complicated mixture that provide a broad array of carbon sources for microbial communities.

**Fig 2.**
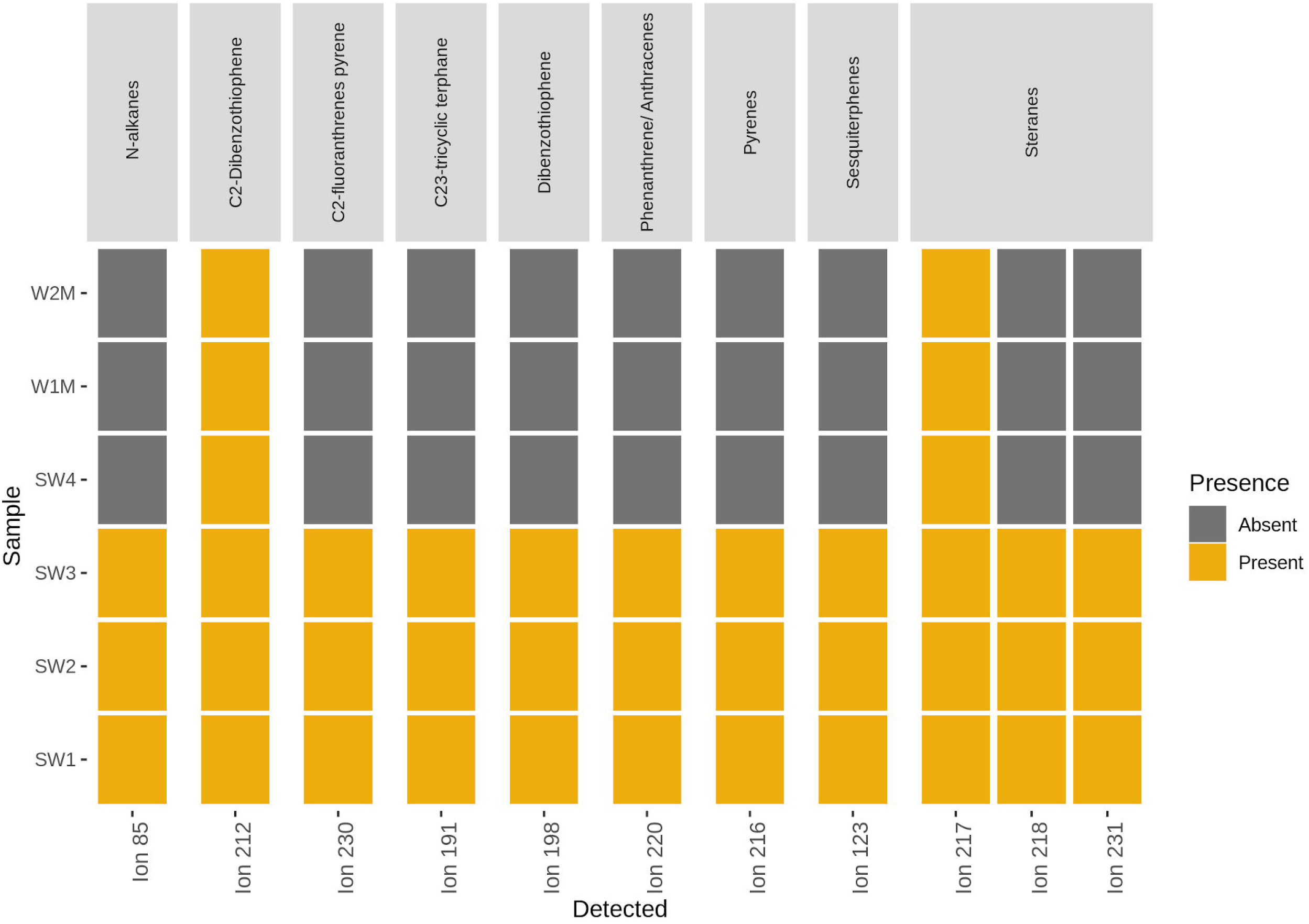
Hydrocarbons detected in the oil well. The thermal map shows the absence (grey) and presence (yellow) of each hydrocarbon analyzed in the water samples. The most representative ion used in the detection of each compound in the mass spectrometer is indicated along the abscissa axis.

### Analysis of microbial communities

The bacterial community in Cahuita N°1 oil well showed a high diversity (Shannon-index > 3) (see Supplementary Fig. S2) and multiple phylotypes. In total, 6993 ASV were identified, of which phylum Proteobacteria was most abundant, representing 60.6 to 79.2 % of the sequences in samples (see Fig. 3a). Within this group Betaproteobacteria represented on average 29.4% of the sequences, Gammaproteobacteria 19.9%, Alphaproteobacteria 20.2 % and Deltaproteobacteria 2.0 %. Other phyla were detected in smaller average proportions, such as Bacteroidetes (4.4 – 11.2 %), Firmicutes (0.5-18.5 %), Candidate Patescibacteria (1.7-15.6 %), Actinobacteria (2.5-6.4 %) and Chloroflexi (0.4-4.6 %) (Fig. 3a).

**Fig 3.**
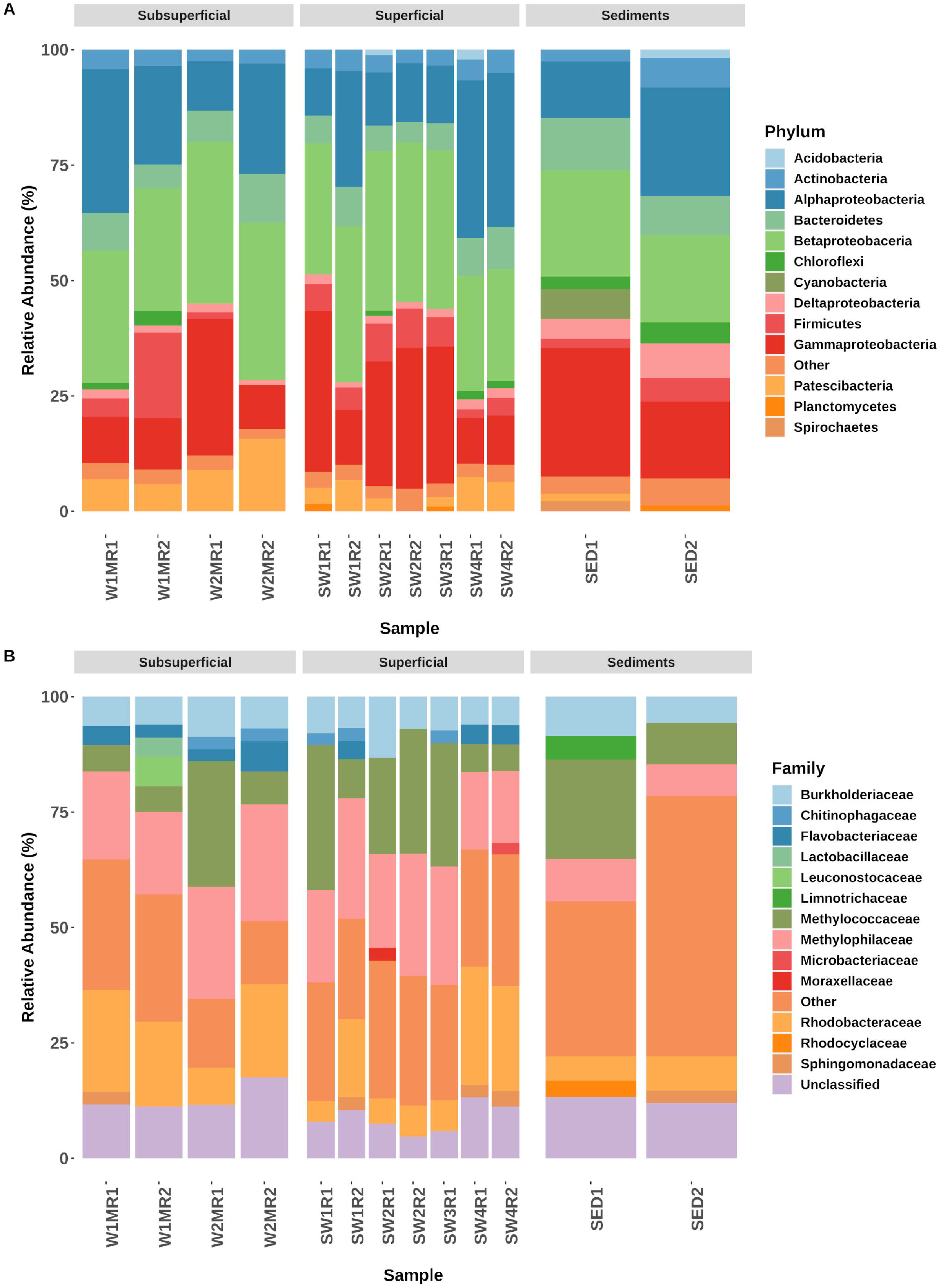
Taxonomic composition of the oil-well waters and contaminated sediments. Relative abundance of bacterial and archaeal organisms at the A) phylum and B) family level. The ASV were taxonomically classified using SILVA reference database v132 [44], as described in Materials and Methods. The superficial water samples are identified as SW1R1 to SW4R2. Sub-superficial water samples are identified as W1MR1 to W2MR2. Sediment samples are identified as SED1 and SED2.

Further analysis of the microbial communities revealed that the dominant families correspond to methylothropic bacteria (Methylophilaceae, Methylococcaceae; 15.7-53.4 %) (see Fig. 3b). Families such as Burkholderiaceae (5.7-13.2 %), Rhodobacteracea (4.5-25.6 %) and Flavobacteraceae (2.6-6.5 %) were also identified with an abundance greater than 2.5 % in more than one sample. These families have been associated with hydrocarbon degradation in contaminated soil and water [13–59–60], governing most of their communities in previous studies. Notably, the family Pseudomonaceae was not an abundant member of the community even though it has been identified as an important hydrocarbon degrader, with a critical role as biosurfactant of hydrocarbons [61].

Abundant ASVs detected in our samples encompass members of the genera *Comamonas*, *Hydrogenophaga*, *Flavobacterium*, *Novosphingobium* and *Terrimonas* (see Fig. 4), bacteria from these taxa have been reported to degrade oil in multiple environments. Particularly they have been related to PAH-contaminated habitats and their biodegradation [62–63], therefore suggesting that they obtain its energy by consuming and degrading the PAHs present in the oil well, ie. dibenzothiophene, phenanthrene, anthracene and pyrene (see Fig. 2). Additionally, previous reports demonstrated that some strains of *Comamonas* and *Hydrogenophaga* can degrade saturated and unsaturated alkanes *in situ* and *ex situ* [62,64]. These ASV have hence likely adapted to the consumption of various hydrocarbons for almost 100 years, making them attractive candidates for isolation. Future work is required to understand the interactions between these bacteria that have co-evolved to degrade hydrocarbons in multiple and complicated sources.

**Fig 4.**
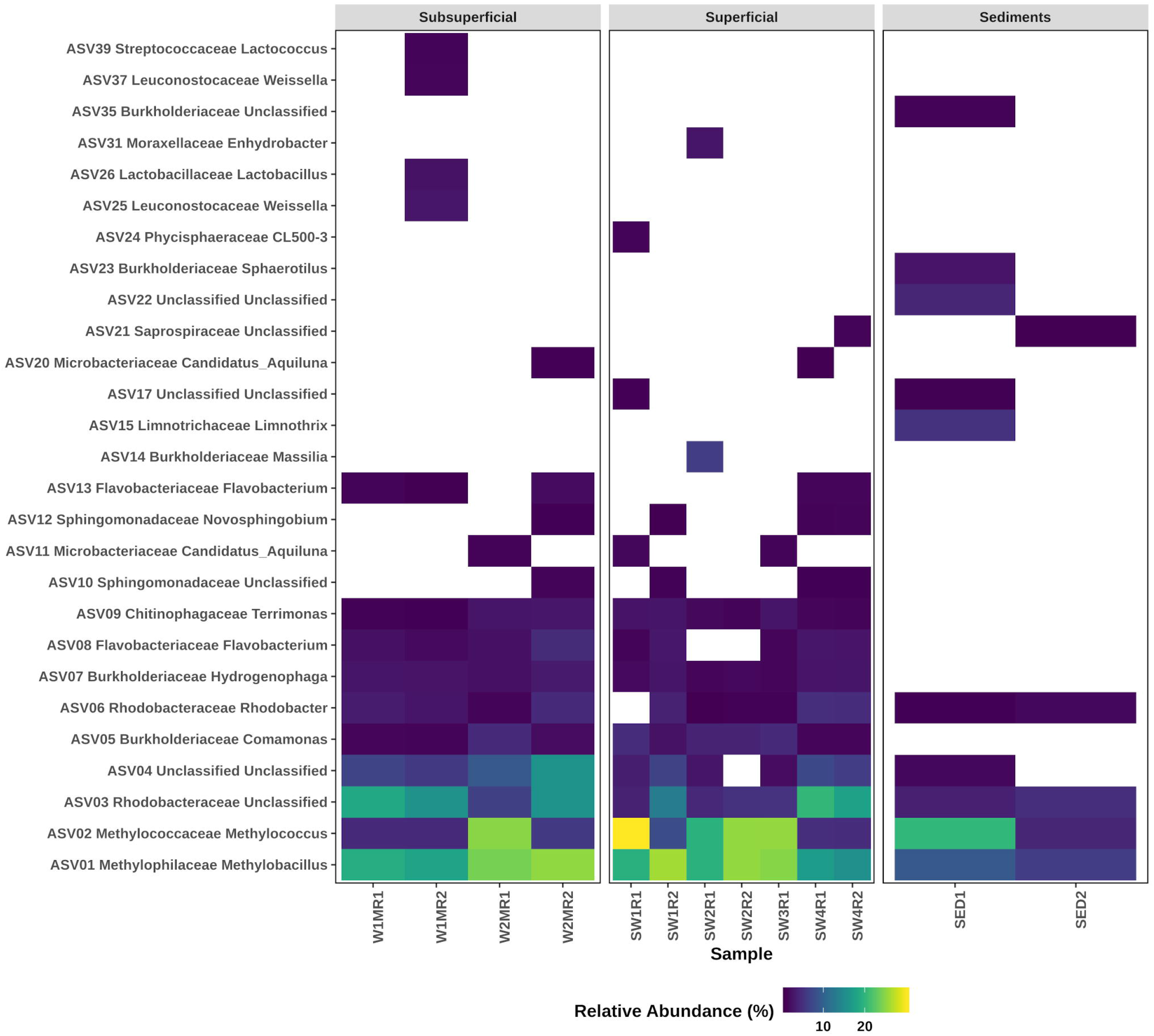
Thermal map representing the most abundant ASV in each sample. This map depicts the percentage of 16S rRNA gene sequences assigned to each ASV (ordinate axis) across the 13 samples analyzed (abscissa axis).

Even though common hydrocarbon consumers were present in the oil well with abundance >1 %, the environment was controlled mainly by C1-consuming bacteria (see Fig. 4), specifically *Methylobacillus*_ASV01 (6.3-26.1 % of total reads) and *Methylococcus*_ASV02 (4.1-30.0 % of total reads). These bacteria are the protagonists in carbon recycling in the trophic chain, performing an essential role in the conversion of CH_4_ to CO_2_. Methylotrophs are characterize for oxidizing C1 compounds such as methane, methanol, and methylamine to the central intermediate formaldehyde. The by-products might be assimilated for anabolism or proceed to the electron chain, consequently, to produce ATP [65]. The ability of these genera to metabolize methane and methanol could hence also participate in the re-coupling of C1 compounds to the trophic chain.

According to our data *Methyloccoccus*_ASV02 is an abundant member of the community (4.1-30.0 %). Members of these genera have been reported to oxidize methane using a soluble or a membrane methane monooxygenase (MMO), an enzyme of which the biological function is to transform methane to methanol [66]. Notwithstanding this effect, this enzyme presents unspecific activity and has the ability to oxidize other hydrocarbons of a longer chain [67]; however, to our knowledge there is no study that demonstrate *Methylococcus* growth with these substrates. The unspecific activity of MMO attributed to *Methyloccocus* and other methane-oxidizing bacteria have an important potential role in the bioremediation of hydrocarbons, specifically short-chain n-alkanes [68–69].

Another important ASV found was *Methylobacillus*_ASV01 which encompass the majority of the sequences classified as these genera. Members of the family Methylophilaceae, where the genus *Methylobacillus* is found, lack the ability to metabolize methane, but they have been reported to be able to use other C1 compounds, such as methanol, methylamine and formaldehyde, as a carbon source [70]. *Methylobacillus* has been found in petroleum-contaminated soils, specifically in experiments of PAH biodegradation in vitro [71], therefore whether they can grow using hydrocarbons as a carbon source is at present unclear. Our chemical analyses revealed that methane is the major gas in the oil well (see Supplementary Fig. S1); other C1 compounds such as methanol were not detected (data not shown; limit of detection for methanol 2.5 mM). We hence suggest that a cross-feeding might occur between different genera such as *Methylococcus*_ASV02 and *Methylobacillus*_ASV01, i.e. *Methylococcus* oxidizes methane to methanol, and *Methylobacillus* uses the methanol that leaks for its growth. The rapid consumption of methanol by these bacteria might explain its absence at levels greater than 2.5 mM in the samples. Similar cross-feeding essential for bacterial survival has been reported in an oil-degrading synthetic consortium, which allows bacteria to survive [72].

Additionally, Non-metric multidimensional scaling (NMDS) and PERMANOVA were consistent in showing significant differences (p = 0.004) between the communities belonging to the superficial water, sub-superficial water and the sediments (Fig. 5). Hydrocarbon-degrading bacteria and their metabolic paths have been shown to differ among aerobic, anaerobic waters and sediments polluted with hydrocarbons [21, 24, 35, 75]. Even though these differences appear in the PERMANOVA, there is no clear difference on analysis of the most abundant groups of the samples (see Fig. 3). The variation in the microbial communities in the water samples might be due to a slight oxygen gradient present between the superficial (0.90 mg O_2_/L, 13.2 % of oxygen saturation) and the sub-superficial samples (0.63 mg O_2_/L corresponding to 6.7 % of oxygen saturation) (see Table 1). This condition might avoid the growth of some microorganisms with metabolisms sensitive to oxygen changes. In contrast, the sediments presented a distinct community because of the volatilization of some hydrocarbons and the accumulation of other oil compounds.

**Fig 5.**
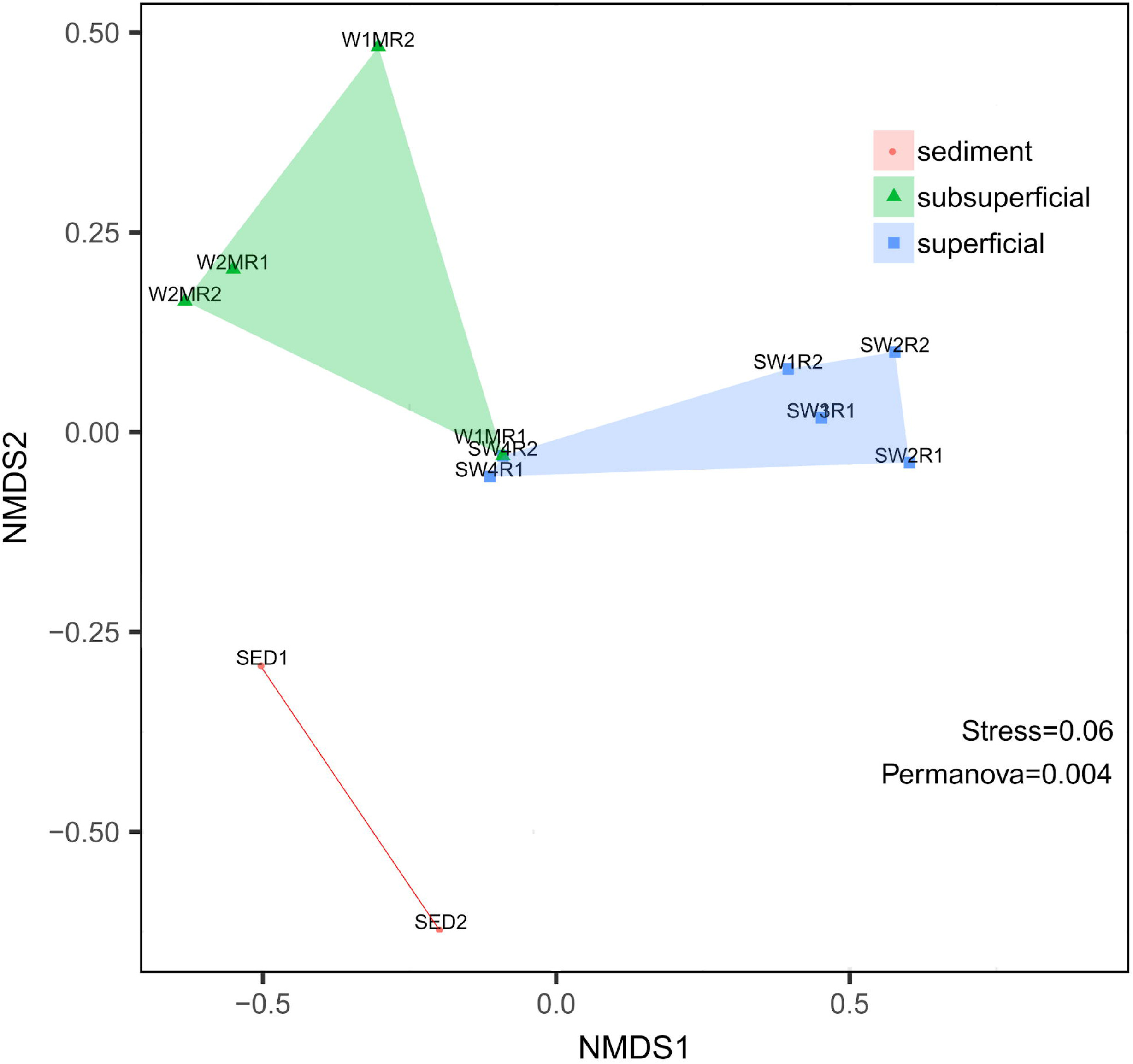
Non-metric multidimensional scaling analysis of the prokaryotic communities in the Cahuita oil well. A clustering of communities according to the oxygen content of the samples (See Table 1) and to the habitat (water column versus sediment) is shown. The NMDS and the Permanova analyses were performed with Package Vegan v2.5-6 [47].

In order to validate the results obtained by 16S rRNA gene amplicons and confirm the importance of the C1 metabolism in this abandoned oil well, we decided to use two approaches: first, identify genes responsible for methane metabolism in the environmental DNA and second isolate methylotrophic bacteria. For the first approximation, we amplify a region of *pmo* gene which encodes for the methane monooxygenase, an enzyme responsible for the oxidation of methane to methanol. As seen in Supplementary Fig. S3, we were able to amplify a band of approximately 700 bp, by performing PCR with degenerate primers previously used to amplify a region of the *pmo* gene. The sequencing of this DNA fragment (GenBank Accession: MT948143) confirms the presence in our samples of a gene that encodes a methane monooxygenase closely related to the enzyme reported in *Methylococcus capsulatus* genome (GenBank Accession: AE017282.2) (93.3% similarity), reaffirming the importance of this genus and its byproducts in the pathways taking place in this abandoned oil well.

On the other hand, for the isolation of methylotrophic bacteria, we try to isolate microorganisms capable of using methane and methanol as the only carbon source. After various efforts, we could not isolate methane-consuming bacteria (data not shown), but we were able to isolate and identify a methanol-consuming bacterium. The strain isolated was a pink-pigmented bacterium which show the ability to use methanol as its sole carbon source in solid and liquid media (See Figs. 6a-6c). According to the 16S rRNA gene sequence the strain was classified as a *Methylorubrum rhodesianum* which is known as a facultative methylotrophic bacterium with the ability to use methanol and methylamine [73]. Phylogenetic analysis revealed that *M. rhodesianum* isolated in this work is closely related to the strain obtained from a formaldehyde enrichment experiment (Fig. 6d) [74]. Interestingly our strain shows a very slow growth in rich media such as R2A and LB (data not shown) suggesting that this strain prefer C1 compounds such as the methanol possibly generated by other bacteria in the oil well. The isolation of bacteria capable of growing solely on methanol reaffirms the importance of C1 compounds in shaping the microbial community in the abandoned oil well. In summary, this system is governed by methylotrophic bacteria and some common hydrocarbon degraders (see Fig. 4), indicating that the community in the oil well is modeled primarily by C1 carbon sources and indicating a community metabolism in which hydrocarbon biodegradation might be coupled with methane metabolism through enzyme specificity or cross-feeding (Fig. 7).

**Fig 6.**
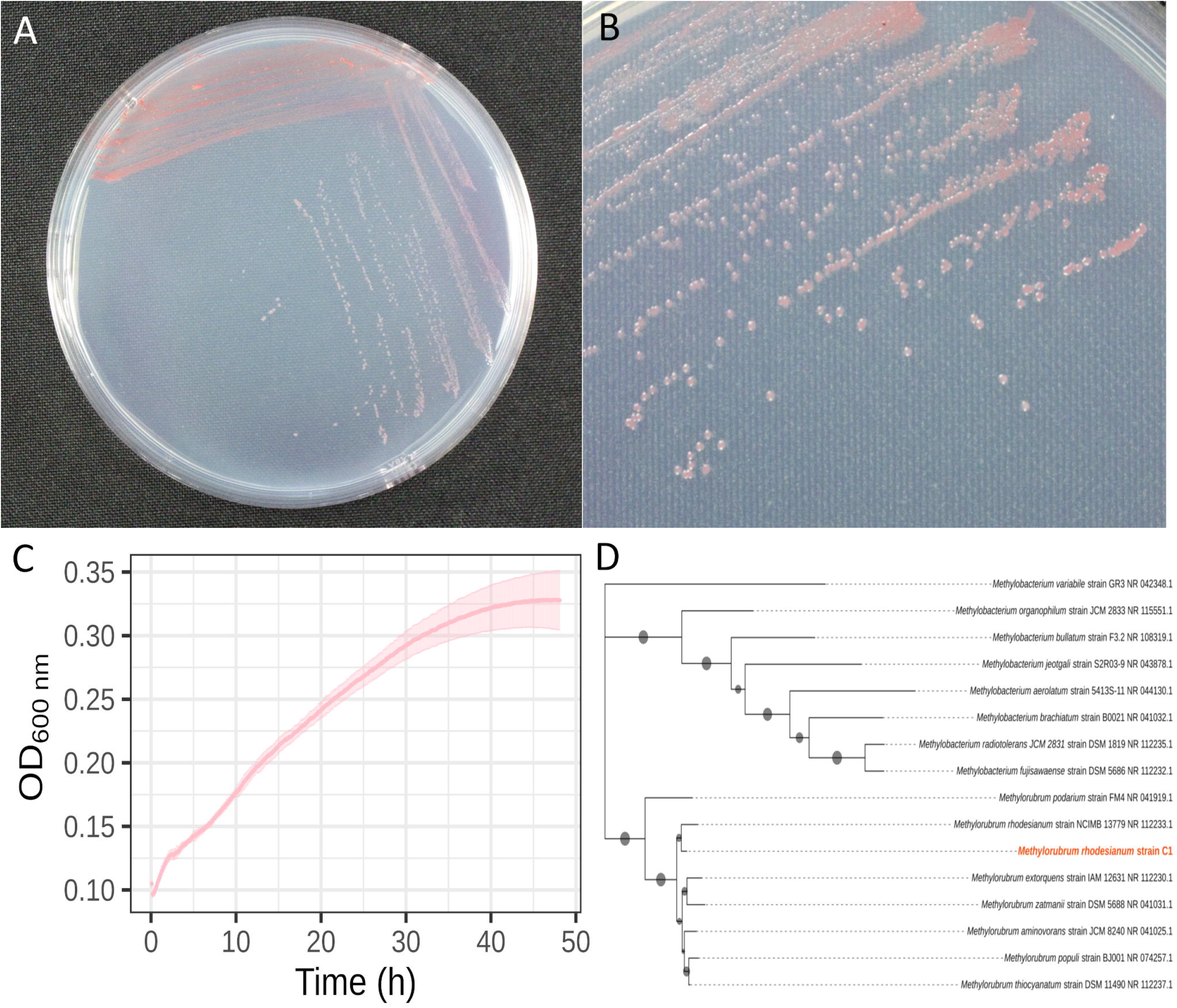
Isolation of *Methylorubrum rhodesianum*, a methylotrophic bacterium present in the abandoned oil well. A) *Methylorubrum rhodesianum* C1 culture in solid M9 mineral medium with methanol (25mM) as its sole carbon source after 48 h. B) Colonies are small and present a characteristic pink color. C) Growth curve of *Methylorubrum rhodesianum* C1 in M9 minimal media with methanol 25 mM as its sole carbon source, the curve was performed as described in material and methods. D) Phylogenetic tree of *Methylorubrum rhodesianum* C1, with close related microorganisms. The phylogenetic reconstruction was made using MEGA as described in Materials and Methods.

**Fig 7.**
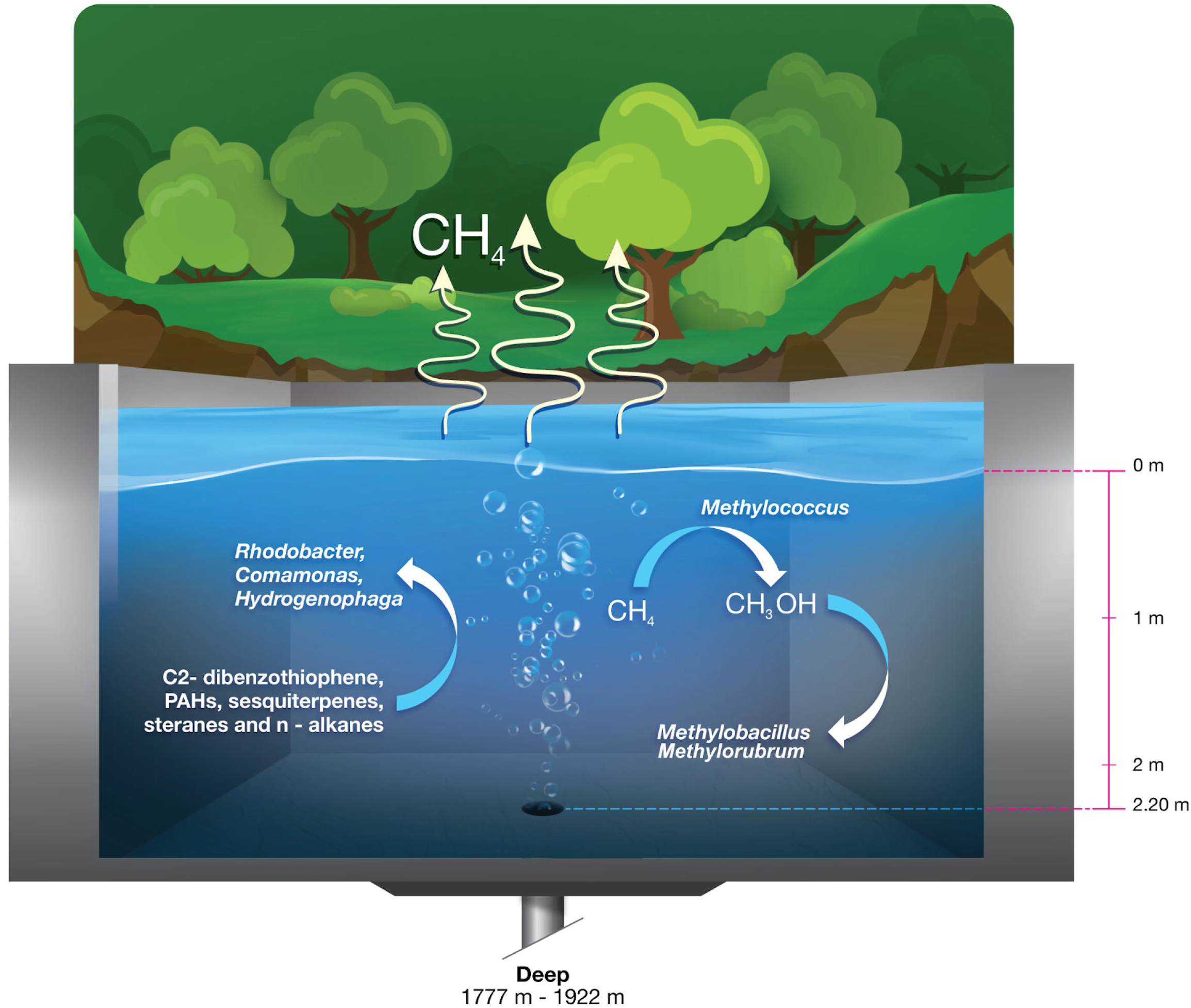
Cartoon of the biochemical processes and microorganisms involved in Cahuita N°1 oil well. The results indicate that the microbial community is dominated by microorganisms capable of using C1 compounds. Methane, possibly of abiotic origin, is oxidized in the first instance by microorganisms of genus *Methylococcus*. The methanol produced can then be used by microorganisms of genus *Methylobacillus* and other facultative methylotrophs such *Methylorubrum*. Other microorganisms capable of using other hydrocarbons were also identified.

## Conclusions

This is the first chemical and microbiological analysis of the abandoned oil well Cahuita N° 1, which has been a focus of pollution in Cahuita National Park for about 99 years. The chemical analysis shows that the Cahuita oil well is characterized by a continuous efflux of methane and the presence of hydrocarbons including C2-dibenzothiophene, phenanthrene or anthracene, fluoranthene pyrene, dibenzothiophene, tricyclic terpene, pyrene, sesquiterpenes, sterane and n-alkanes in a complicated mixture. The microbial community in the oil well is dominated by methylotrophic bacteria (e.g. *Methylobacillus*, *Methylococcus* and *Methylorubrum*) and other common hydrocarbon degraders (*Burkholderia*, *Rhodobacter* and *Flavobacterium*). Our results show that the community is governed by a C1 metabolism, conditioned by the methane expelled by the oil well (Fig. 7). The enduring contamination makes this habitat of great interest for bioremediation purposes, including the isolation of novel hydrocarbon-degrading bacteria long-term adapted to complex mixtures of these compounds. Future work is required to obtain more bacterial isolates and to understand the whole biochemical reactions occurring in this abandoned oil well.

## Acknowledgements

We are grateful for the support of park rangers at Cahuita National Park and the SINAC administration and to Arnoldo Vargas and José Jimenez for assistance in the design of some figures.

## Author contributions

CCR and MC conceived and designed the experiments; DRG, RA, RAl, PF, DP-P performed the experiments; DRG, PF, MC, KR analyzed the data; PF, DP-P MC contributed reagents or materials or analytical tools; DRG, PF, KR, MC wrote the paper. All authors reviewed and approved the final version of the manuscript.

## Availability of data and material

All sequences obtained in this study are available at BioProject ID PRJNA614582 and GeneBank accession numbers MT948143 and MT936438.

