## Supplementary information for "C1 compounds shape the microbial community of an abandoned century-old oil exploration well"

^1^Centro Nacional de Innovaciones Biotecnológicas (CENIBiot), CeNAT-CONARE, 1174-1200 San José (Costa Rica). ^2^Centro de Investigación en electroquímica y Energía química (CELEQ), Universidad de Costa Rica, 11501-2060 San José (Costa Rica). ^3^Escuela de Química, Universidad de Costa Rica, 11501-2060 San José (Costa Rica). ^4^Escuela de Biología, Universidad de Costa Rica, 11501-2060 San José (Costa Rica). ^5^Programa Institucional de Fomento a la Investigación, Desarrollo, e Innovación, Universidad Tecnológica Metropolitana, Santiago, Chile ^6^Hidro Ambiente Consultores, 202, San José (Costa Rica), ^7^Centro de Investigaciones en Productos Naturales (CIPRONA), Universidad de Costa Rica, 11501-2060 San José (Costa Rica).

Keywords: Methylotrophic bacteria, Methylobacillus, Methylococcus, Methylorubrum, Hydrocarbons, Oil well, Methane, Cahuita National Park

^*^Correspondence to: Max Chavarría

Escuela de Química & Centro de Investigaciones en Productos Naturales (CIPRONA)

Universidad de Costa Rica

Sede Central, San Pedro de Montes de Oca

San José, 11501-2060, Costa Rica

Phone (+506) 2511 8520.  Fax (+506) 2253 5020

###

**SUPPLEMENTARY TABLES**

**Table S1. DNA sequence and phylogenetic assignment of most abundant phylotypes detected in Cahuita exploratory oil well N^o^ 1 using Illumina-based amplicon deep-sequencing.**

See Excel file.

**LEGENDS OF SUPPLEMENTARY FIGURES**

**Fig. S1 Diversity measures of the samples in Cahuita exploratory oil well N^o^ 1**. The diversity measures (Shannon, Simpson and Observed Richness) were calculated using phyloseq. Figure shows A) diversity measures of all samples grouped by sample point. B) diversity measures of all samples.

**Fig. S2 Chromatogram of the gas sample obtained from Cahuita exploratory oil well N^o^ 1**. The figure shows chromatograms of the gas sample from the oil exploration well (above) compared with a methane standard (below).

**Fig. S3 Amplification of *pmo* genes in** **Cahuita abandoned oil well samples**. To evaluate the presence of the *pmo* gene in our samples, we performed a PCR as described in Materials and Methods section. The results revealed the presence of the *pmo* genes in Cahuita abandoned oil well samples (lane 2-9). A negative control was prepared with all components of the PCR reaction except a DNA sample (lane 10). For further confirmation of the gene identity sequencing of the SW2 was performed.

**LEGEND OF SUPPLEMENTARY VIDEO**

**Video S1. Gas efflux in Cahuita exploratory oil well N^o^ 1, Limon, Costa Rica.**
