## Supplementary figures and images for "C1 compounds shape the microbial community of an abandoned century-old oil exploration well"

### Figure S1

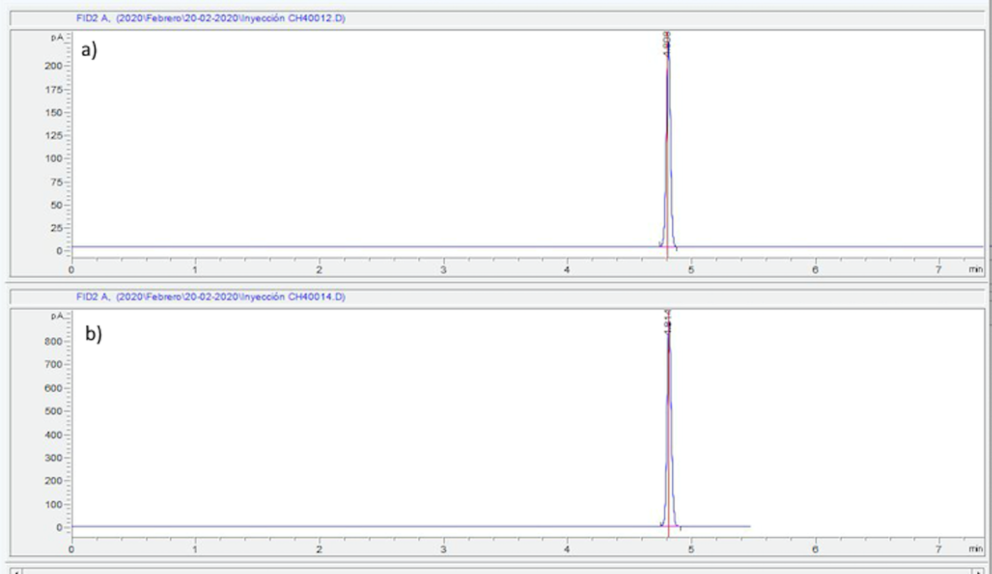

### Figure S2

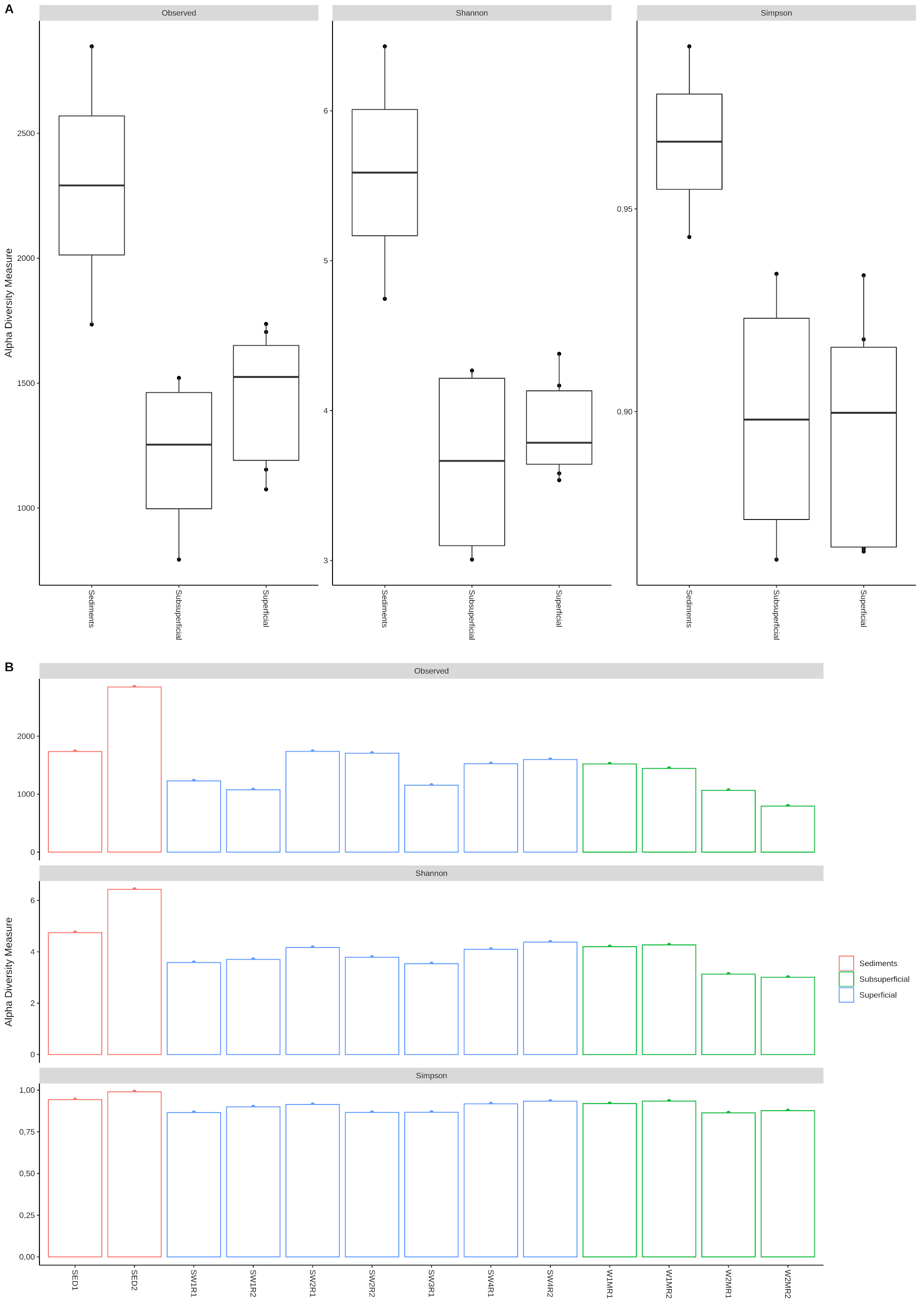

### Figure S3

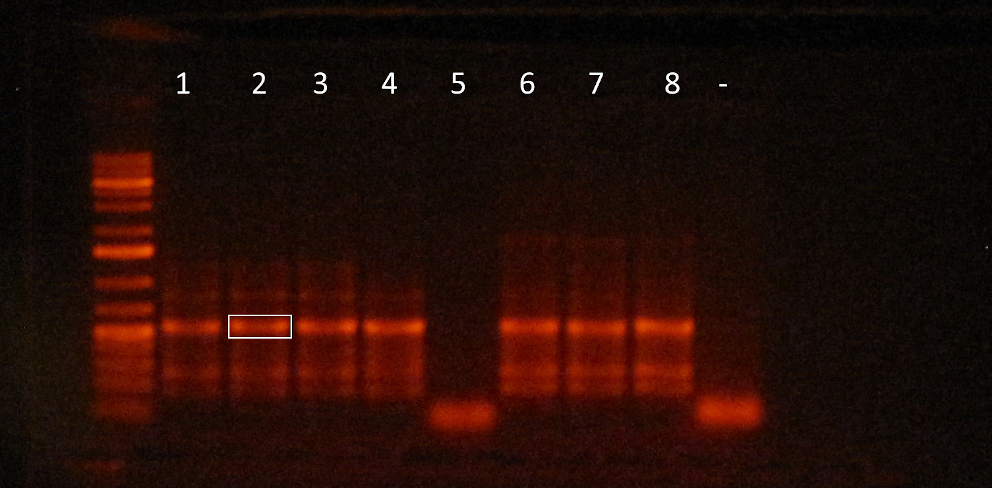
